# Structural Analysis of Prostate Cancer N-Glycans Using Graph-Based Structural Metrics

**DOI:** 10.64898/2026.06.12.731995

**Authors:** Mihir Kalyanthaya, Daniel Gallegos, Joleen Chow, Bruno Pavletic, Aurora Britania Diaz Fernandez, Michelle Kilcoyne, Lokesh Joshi, David H. Nguyen

**Author notes:** Corresponding author: David H. Nguyen.

## Abstract

The N-linked glycans are structurally complex carbohydrate modifications that regulate protein folding, immune recognition, and cellular signaling, and their expression is extensively remodeled during cancer progression, making them promising biomarkers. In this study, prostate cancer–associated N-glycans from a range of relevant peer-reviewed studies were curated and digitized to develop a versatile computational framework that quantitatively encodes their spatial complexity across diverse biological systems. We invented two indices—the Distance & Connectivity Index (DCI) and the Position & Composition Index (PCI)—to capture the spatial information in N-glycans as layered architectures, enabling calculation of residue-level path lengths, branching structure, and compositional diversity. DCI summarizes glycan structure as both a scalar and matrix representation, while PCI does the same but also captures monosaccharide diversity, linkage heterogeneity, and cross-layer branching features. These metrics were computed with GlycoAssessor, an open-source platform that extracts information for the DCI and PCI from glycans drawn via Symbol Nomenclature for Glycans (SNFG) notation. Principal Component Analysis (PCA) was applied to evaluate whether glycans from prostate cancer tissues cluster distinctly in a disease-relevant manner. Results show that the spatial information in N-glycans: (1) increased in a multi-dimensional, non-linear manner, (2) objectively segregated structural themes, (3) could function as a potential prostate cancer biomarker that is distinct from mass-to-charge ratio and relative abundance, and (4) could objectively quantify novel subtype classifications of glycans associated with disease states and progression.

## Introduction

Protein glycosylation is among the most prevalent and structurally diverse post-translational modifications in eukaryotic cells. By covalently attaching carbohydrate chains to proteins, glycosylation influences protein folding, stability, trafficking, receptor activation, and immune recognition (Tian et al., 2025). Unlike nucleic acids and proteins, glycan biosynthesis is not template-driven, resulting in a high degree of structural heterogeneity that is tightly regulated by cellular and molecular context. This inherent complexity allows glycans to function as a rich informational layer in biological systems to promote cell signaling and differentiation but also presents significant analytical challenges (Li et al., 2024).

Among the various forms of glycosylation, N-linked and O-linked glycans represent the two dominant classes of protein-associated glycans in mammals. Their distinct biosynthetic pathways and structural features give rise to different functional roles in health and disease. N-glycans are covalently attached to the amide nitrogen of asparagine residues within the conserved Asn-X-Ser/Thr consensus sequence. Their biosynthesis initiates in the endoplasmic reticulum (ER) with the en bloc transfer of a preassembled oligosaccharide to the nascent polypeptide. Subsequent trimming and remodeling in the ER and Golgi apparatus generate a diverse repertoire of mature N-glycan structures (Hang et al., 2015). Despite this diversity, all N-glycans share an invariant pentasaccharide core (Man GlcNAc) consisting of three mannose (Man) monomers conjugated to a chitobiose core comprised of two N-acetylglucosamine (GlcNac) residues, which serves as the structural foundation for branching and elongation. N-glycans play essential roles in protein quality control, cell signaling, immune modulation, and intercellular communication. Dysregulation of N-glycan biosynthesis has been implicated in inflammation, cancer, and congenital disorders of glycosylation (Reily et al., 2019; Freeze et al., 2009; Pandey et al., 2022).

O-glycans, in contrast, are attached to the hydroxyl oxygen of serine or threonine residues and do not arise from a universal precursor. O-glycosylation typically begins in the Golgi with the addition of a single monosaccharide, most commonly N-acetylgalactosamine (GalNAc) in mucin-type O-glycans, followed by stepwise elongation. O-glycans are especially abundant on mucins and cell-surface proteins, where they contribute to lubrication, pathogen defense, immune evasion, and metastatic behavior (Wei et al., 2026; Chia et al., 2016; Zhu et al., 2023; Tsuboi et al., 2012).

Despite their heterogeneity, mammalian N-glycans exhibit conserved structural principles shaped by shared biosynthetic pathways. The invariant Man GlcNAc core from N-glycans forms a basis for three canonical structural classes: high-mannose, hybrid, and complex (Alley et al., 2013). Branching architecture is a defining feature of complex N-glycans. Biantennary, triantennary, and tetraantennary configurations correspond to increasing numbers of glycan arms extending from the core. The degree of branching is regulated by N-acetylglucosaminyltransferase activity and has been linked to cellular differentiation, immune regulation, and oncogenic transformation (Mehta et al., 2012; Lau et al., 2007). Additional modifications, such as core fucosylation, terminal sialylation, bisecting GlcNAc, and polylactosamine repeats, further modulate glycan function by influencing charge, receptor binding, and immunogenicity (Shirakawa et al., 2021).

Cancer cells undergo widespread remodeling of protein glycosylation as a consequence of altered metabolic state, Golgi organization, and glycosyltransferase expression (Reily et al., 2019) . Several recurring N-glycan alterations have been observed across diverse tumor types. One of the most prominent is increased β-(1,6)-linked GlcNAc branching, driven by upregulation of N-acetylglucosaminyltransferases, which enhances cell migration, invasion, and growth factor receptor signaling (Mehta et al., 2012). Increased sialylation is another common feature, contributing to immune evasion, resistance to apoptosis, and metastatic potential by masking cell-surface antigens and altering receptor interactions (Vasudevan & Haltiwanger, 2014). Altered fucosylation of N-linked glycans, including increased core fucosylation and aberrant outer-arm fucosylation, modulate cell–cell adhesion, receptor signaling, and antibody interactions have emerged as important cancer biomarkers (Keeley et al., 2019; Miyoshi et al., 2012). Additionally, some tumors exhibit increased levels of high-mannose or immature N-glycans, reflecting dysregulated Golgi processing or cellular stress responses (de Leoz et al., 2010; Oh et al., 2022). Collectively, these alterations underscore the diagnostic and therapeutic (Khan et al., 2026) potential of cancer-associated glycan signatures.

Prostate cancer exhibits a distinct and well-characterized set of N-glycan alterations that correlate with tumor grade, metastatic potential, and disease recurrence. Increased core α-(1,6)-linked fucosylation is frequently observed in prostate tumors and has been linked to altered androgen receptor signaling and the development of aggressive, castration-resistant disease (Höti et al., 2018; Ippolito et al., 2024). Increased branching, particularly β-(1,6)-branched bi-, tri-, and tetraantennary structures, is associated with enhanced tumor cell survival, invasiveness, and growth factor signaling (Ishibashi et al., 2014; de-Souza-Ferreira et al., 2023). Prostate cancer cells also display elevated sialylation, which contributes to immune evasion and modulation of metastatic behavior. Notably, increased retention of high-mannose N-glycans has been observed in prostate tumors and metastases, suggesting dysregulated Golgi processing and altered receptor trafficking (Kapp et al., 2025). Changes in the N-glycosylation of prostate-specific antigen (PSA), including increased core fucosylation and sialylation, have been shown to improve differentiation between benign and malignant conditions, highlighting the clinical relevance of glycan structural changes (Saldova et al., 2010; Llop et al., 2016; Kailemia et al., 2016).

Several standardized representation systems have been developed for glycans, including IUPAC, GlycoCT (Herget et al., 2008), and WURCS (Tanaka et al., 2014). IUPAC notation provides a human-readable format that explicitly names monosaccharides and linkage types, making it useful for communication and visualization (McNaught, 1996; Varki et al., 2015). GlycoCT offers a highly structured, machine-readable format that encodes stereochemistry and linkage positions and is widely used in glycoinformatics databases (Herget et al., 2008; Tsuchiya et al., 2018). WURCS provides a canonical, database-friendly identifier capable of representing extremely complex and ambiguous glycans, albeit at the expense of human readability (Tanaka et al., 2014; Matsubara et al., 2017). While these languages excel at encoding chemical identity, they were not designed to capture higher-order spatial features such as branching patterns, hierarchical depth, or connectivity patterns. As a result, they are limited in their ability to support quantitative comparisons based on structural geometry. The difference between the purpose of the DCI and PCI compared to these existing languages is to quantitatively capture the structural information in glycans, not to translate it.

This study introduces the Distance & Connectivity Index (DCI) and the Position & Composition Index (PCI) as algorithms that objectively interpret the spatial information in N-glycans. We validate the DCI and PCI as generalizable algorithms for classifying N-glycan structural themes and distinguishing glycan subtypes using spatial information alone. We demonstrate that the spatial information in N-glycans reveal objective, quantitative insights in the form of unknown prostate cancer subtypes based on N-glycosylation hidden to mass-to-charge (m/z) ratios and the visual classification of glycan structure. By demonstrating that glycan spatial information encodes disease-relevant states, this framework furthers glycomics toward quantitative spatial analytics and introduces a new method for potential biomarker discovery. Ultimately, identifying these spatial glycan signatures not only uncovers hidden disease states and stages of progression, but also opens new translational pathways for targeted therapies and improved patient diagnostics.

## Methods

The Distance & Connectivity Index (DCI) and the Position & Composition Index (PCI) represent glycans as mathematical graphs rather than chemical entities. First, in the PCI framework, monosaccharides are treated as nodes, glycosidic linkages as edges, specific anomeric configurations (alpha or beta) and carbon positions extracted from the SNFG drawings enable the application of graph-theoretic concepts. The PCI decomposes the glycan into hierarchical layers to compute numerical descriptors related to residue counts, linkage diversity, and inter-layer connections. Second, in the DCI framework, while monosaccharides are treated as nodes and glycosidic linkages as edges, the nature of the glycosidic linkage is ignored; the DCI only quantifies features such as path length from the root and branching complexity. Both the PCI and DCI produce a matrix representation and a scalar value summarizing overall connectivity.

### The Position & Composition Index

To calculate the PCI, the glycan is first divided into layers, starting from the base node that is attached to the asparagine residue. Layers are defined as monosaccharides separated by one linkage bond (Figure 1A). Within a layer, a monomer may not be connected to any other monomers of that layer. Therefore, connected units must be in adjacent, but vertically separate layers. A fucose that is attached to the base node is counted as belonging to the next level up the chain since a layer is defined by being separated by one glycosidic linkage. Once the monosaccharides have been assigned to their layers, the following five features are counted for each layer.

1. <u>N</u>umber of <u>M</u>onosaccharide <u>T</u>ypes (NMT) - This counts how many unique monosaccharide types are in a layer.
2. <u>T</u>otal <u>N</u>umber of <u>U</u>nits (TNU) - This counts the total number of monosaccharides of any type in a layer.
3. <u>N</u>umber of Int<u>e</u>r-layer <u>L</u>inkages (NEL) - This counts the total number of glycosidic linkages that connect a layer with the layer above it *and* the layer below it.
4. <u>N</u>umber of <u>L</u>inkage <u>T</u>ypes (NLT) - This counts only the unique types of glycosidic linkages, ignoring replicates, that separate a layer from the layer above it *and* the layer below it.
5. <u>N</u>umber of <u>M</u>onomers <u>C</u>onnected to a <u>D</u>ifferent <u>S</u>pecies in the <u>P</u>revious <u>L</u>ayer (NMCDSPL) - This counts the number of monomers in a layer that are connected to a different monosaccharide type in the previous layer.

**Figure 1.**
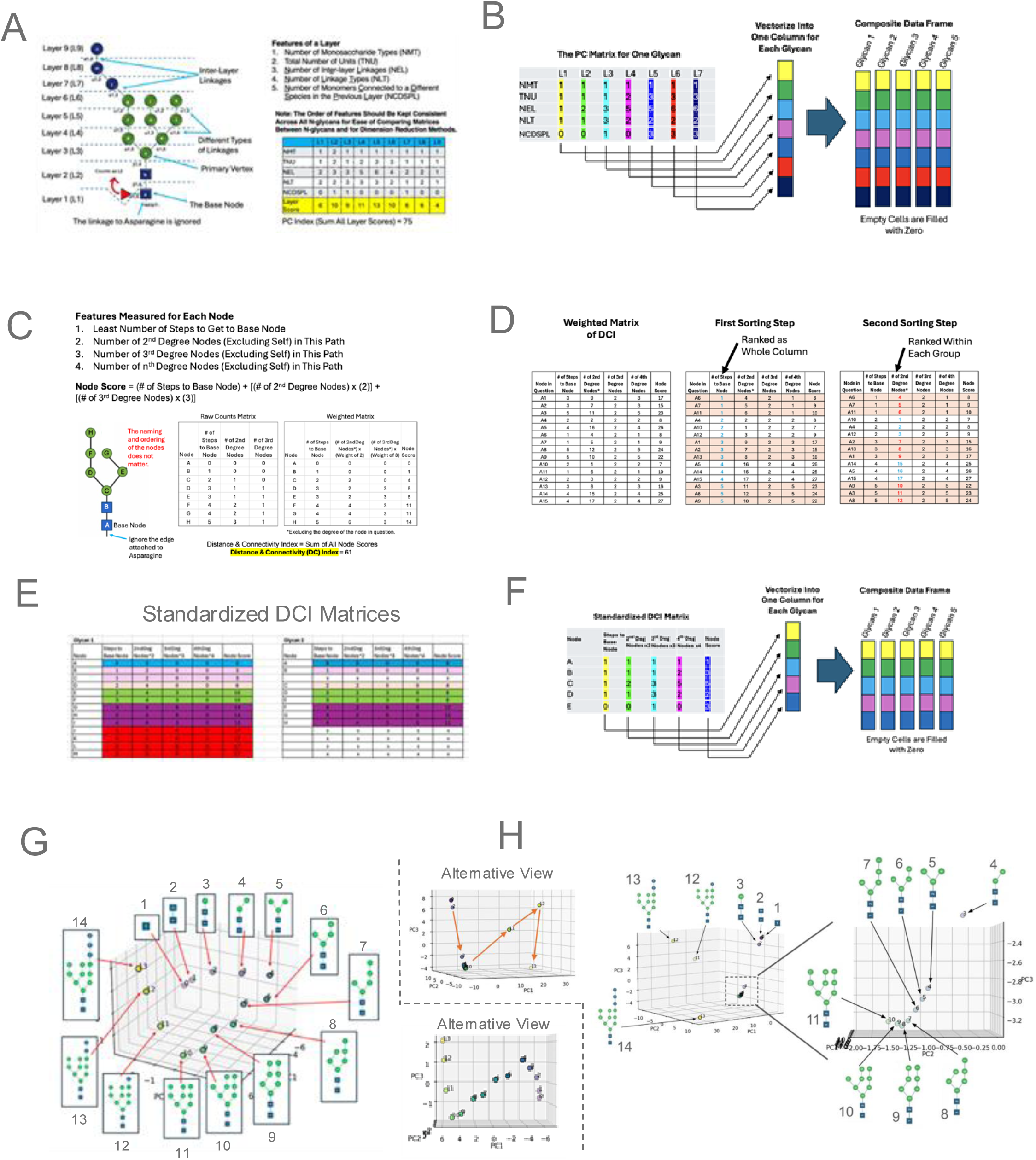
How the Position and Composition Index (PCI) and the Distance and Connectivity Index (DCI) Are Derived. (A) The PCI calculation diagram. The glycan is divided into unique layers (L1 means “Layer 1”) starting from the base, where a layer is defined by the distance of one linkage. Thus, a fucosylation is included in the next layer up. PCI features are defined above the matrix used to calculate the scalar index. (B) PCI vectorization process. Each layer in the PC matrix is stacked (color coded for clarity) to create a single column. This is repeated for each glycan in a data set. Systematic vectorization allows for meaningful PCA analysis. Any empty cells are filled with zero. A Python script that automates this process is available (Nguyen, 2026a). (C) DCI calculation diagram. The glycan is labeled per sugar starting from the base. The exact names of each node does not matter. DCI features are defined along with the formula for calculating a node score. Counting features results in a “raw counts matrix” that is converted to a “weighted matrix” that contains a column of node scores that can be summed into a scalar index. (D) Column sorting of the DCI matrix in preparation for dimension reduction techniques. Two sorting steps are required. First is to sort the column called “# of Steps to Base Node”, which ranks each node from least to greatest. Second is to identify unique subsets based on the previously sorted column and then sort each subset according to the column called “# of 2nd Degree Nodes”. (E) DCI matrix standardization. Glycans must be standardized by adding empty cells with a ‘0’ (in the diagram, the 0 are represented by ‘x’) to ensure that all the glycans align for meaningful dimension reduction analysis. (F) The vectorization process of a standardized DCI matrix. Each layer in the standardized DCI matrix is stacked (color coded for clarity) to create a single column. This is repeated for each glycan in a data set. A Python script that automates this process is available (Nguyen, 2026b). (G, H) The 14 canonical eukaryotic N-glycans analyzed by the PCI (G) or DCI (H). The first three principal components are plotted. Each data point is labeled with the structure and step number. The alternative view shows non-linearity.

Figure 1A shows the resulting data matrix from counting the above five features for each layer, starting with the first layer, and arranging the results as consecutive columns from left to right. Each column of values is summed to derive a layer score. Then all layer scores can be summed to derive a total score that is the DCI. While the DCI is a convenient scalar value, the DCI matrix from which it was derived harbors a lot of valuable spatial information that is lost when collapsed into one value. Thus, a systematic way of vectorizing the DCI matrix of a glycan into a one-dimensional array is necessary to compare the DCI matrices of multiple glycans via a dimension reduction technique.

### Systematic Vectorization of the Position & Composition Index for Dimension Reduction

While a scalar index is simple to comprehend, the true richness of spatial information obtained via the PCI is found in the matrix from which the scalar index is derived, referred to as the PCI matrix. Thus, in order to access the multidimensional information in the matrix there needs to be a way to systematically turn each m x n matrix into an m x 1 vector, otherwise known as a column. In the form of a vector, many glycans can be combined as columns in a data frame on which dimension reduction techniques, such as Principal Component Analysis (PCA), can be applied. However, in order for the results of dimension reduction to be meaningful, the rows of each column must represent the equivalent feature. Thus, a vectorization protocol is required that objectively and systematically turns each PCI matrix into a vector that has features ordered in a manner that matches all glycans being compared as a set. Figure 1B exhibits how each column in a PCI matrix, representing each layer of the glycan as defined by the PCI, is subsequently stacked under the previous layer, starting from Layer 1. In each layer, the following order of features is always maintained: NMT, TNU, NEL, NLT, and NCDSPL. Thus, for each layer in the vectorized matrix, this order is repeated. For matrices of glycans that have fewer layers than the glycan that has the most layers, empty cells are filled with a zero. In this way, the PCA space that results from the composite dataframe of many vectorized glycans is not universal, but always relative to the glycan in the set that has the most layers. A Python script that automates this vectorization process for PCI matrices is available on GitHub (Nguyen, 2026a).

### The Distance & Connectivity Index

The DCI was designed to capture the complexity of connectivity in N-glycans, especially since they exhibit recurring motifs of bifurcations and trifurcations at specific junctions. As 3D molecules, a bifurcation or trifurcation created by the attachment of a new monosaccharide would induce steric rearrangements that change the shape of the glycan. Thus, being able to quantify the degree of branching behavior may reveal spatial insights that are distinct from what the PCI captures.

The DCI treats glycans as a system of nodes and edges, with the monosaccharides being nodes and the linkages being the edges. The types of sugars and linkages don’t matter, nor do the names of the nodes. The first node is named starting with the letter A, followed by B, then C, and so on and so forth. Naming the sugars in this way is meant to create a convenient way of identifying each sugar but has no effect on the results. As shown in Figure 1C, the first feature of interest is the distance of each node to the base node. The base node is defined as the first monosaccharide that is attached to the asparagine residue. Distance is defined as the least number of steps to get to the base node. The second to fourth features of interest that will be described next involve “degrees”. The degree of a node is defined by how many other nodes it connects with: for example, a node is 2nd degree if it connects with two other nodes. It is important to note that when considering whether a node is 2nd, 3rd, or 4th degree, the node itself is excluded. This is to prevent a node from connecting to itself, as in an edge that extends from a node and then turns around to connect back to the same node. Self-connecting nodes are possible in the graph theory concepts that were adapted for this study, but are irrelevant to glycan structures. The second feature of interest in the DCI is the number of 2nd degree nodes that occur along the path of getting to the base node. The third feature is the number of 3rd degree nodes along the path of getting to the base node. Lastly, the fourth feature is the number of 4th degree nodes along the path of getting to the base node – the highest possible degree of a node is 4th. These four features comprise what is referred to as the “raw counts matrix” (Figure 1C, Step 2).

In order to emphasize the impact of having 2nd, 3rd, and 4th degree nodes, a new matrix is derived by multiplying weights to certain columns of the raw counts matrix. Figure 1C Step 3 shows that the “Weighted Matrix” results from multiplying each 2nd degree node with a weight of 2, each 3rd degree node with a weight of 3, and, if existent, each 4th degree node with a weight of 4. For each node, represented by a row in the Weighted Matrix, a Node Score is obtained by summing the values in that row. Summing all values in the Node Score column then results in a scalar value representing the DCI of a glycan.

### Systematic Vectorization of the Distance & Connectivity Index for Dimension Reduction

Similar to PCI vectorization, vectorizing the DCI allows access to the multidimensional richness of the DCI, which is hidden when reduced to a scalar index. For the DCI matrix, vectorization is more complex but the same constraints apply in order for the results of dimension reduction are meaningful: (1) the order of each feature in the vector must be the same across all glycans that are converted to a vector for comparison as a set of glycans; (2) every vector in a set of glycans being compared must have the same number of rows. To meet these constraints, two sorting steps must occur. Figure 1D shows that the first sorting step is to sort the column representing “# of Steps to the Base Node” in the Weighted Matrix by ascending order. Then, every row that has the same value in “# of Steps to the Base Node” must be grouped as a subset. Once all unique subsets are identified, the second sorting step is applied to each of them separately, but this time sorting by the column representing “# of 2nd Degree Nodes” in ascending order. These first two steps result in an objective and systematic way of sorting all Weighted Matrices from the DCI. However, more complicated N-glycans will have a larger matrix than simpler ones and vectorizing these matrices would result in vectors of unequal lengths and incongruent row identities. Thus, for any set of glycans being compared via dimension reduction, their Weighted Matrices must be standardized to have the exact same m x n dimensions. Figure 1E shows that this is accomplished by ensuring that every glycan being compared must have (a) the exact same number of subsets defined in Figure 1D and (b) the exact same number of rows within each subset. New rows are filled with zeros, represented by “x” in Figure 1E, resulting in what is referred to as a Standardized DCI Matrix. This step ensures that all glycans being compared via dimension reduction on the DCI matrix have the same m x n dimensions. The implication of this step is that the PCA space, assuming PCA is applied to a set of vectorized matrices, is not absolute but always relative to the most complicated glycan in the set. Note that though the most complex glycans in a set of glycans being compared have the most influence on determining the dimensions of a Standardized DCI Matrix, simpler glycans can have rows that do not exist in more complex glycans. Once in their standardized form, these matrices are ready for vectorization by stacking each column under the previous column, starting from the first column, which is “# of Steps to the Base Node” (Figure 1F). Multiple vectors can then be combined as columns in a data frame upon which dimension reduction is applied. A Python script that automates this vectorization process for DCI matrices is available on GitHub (Nguyen, 2026b).

### GlycoAssessor

GlycoAssessor is a desktop application built using the Python (3.14.4) Tkinter GUI toolkit. The open-source scripts are available on GitHub; the ReadMe file has instructions for finding the subfolders that contain the installation files for Windows or Mac operating systems (Gallegos, 2025a). Video tutorials for installing GlycoAssessor are available for Mac computers (Gallegos, 2025b) and Windows computers (Kalyanthaya, 2025a). A video tutorial on how to use GlycoAssessor is also available (Kalyanthaya, 2025b). GlycoAssessor allows the user to draw glycans by connecting individual monosaccharide nodes with α- and β-glycosidic linkages in a two-dimensional canvas, and to automatically calculate DCI and PCI matrices from the drawn glycan. The seventy-five monosaccharides included in the application follow the Symbol Nomenclature for Glycans (SNFG) (Neelamegham et al., 2019). Index and matrix values are calculated by converting the glycan into Python dictionaries of nodes and linkages, computing the path to the base node, then storing the relevant attributes of each node and layer in a Pandas data frame. This data frame is printed to the display log and saved in CSV format.

### Numerical Methods

PCA was done in the Python language using the SciKit Learn library. Vectorized matrices were combined into one data frame and median-centered before doing PCA. The scripts for vectorizing PCI matrices (Nguyen, 2026a) and the DCI matrices (Nguyen, 2026b) were written in Python and available on GitHub. Video tutorials explaining how to run the Jupyter Notebooks are linked to in their associated ReadMe files on GitHub.

## Results

To prove the efficacy of the PCI and DCI in extracting unique spatial information from the structure of N-glycans, they were applied to glycans from various published studies. In each case, they revealed biological insights not attainable via m/z ratio and relative abundance.

### The Spatial Information in N-Glycans Accrues Non-Linearly

The canonical ER synthesis pathway of N-glycans in mammals (Figure S1) begins with the addition of a GlcNAc residue to a dolichol-pyrophosphate (Stanley et al., 2022). The sequential addition of 13 more monosaccharides results in the mature N-glycan precursor Glc_3_Man_9_GlcNAc_2_-P-P-Dol. As an example of how the DCI and PCI interprets the structural information in N-glycans, both were applied to these 14 canonical structures. Plotting the first three principal components of PCA of the vectorized PCI matrix (Figure 1G) or DCI matrix (Figure 1H) revealed that the accrual of spatial information in this pathway increases non-linearly. In fact, it changes in an elegant sinusoidal swirling pattern, highlighting the complex but quantitatively systematic way in which information is stored in N-glycan structure. While encodings like IUPAC, GlycoCT, and WURCS are translations of glycan structure, the PCI and DCI algorithms are more like interpretations of glycan structure, because it is not possible to reconstruct the exact N-glycan by going in reverse from a DCI or PCI matrix. Of note is the difference between what the PCI and DCI captures. The 14 structures of the canonical pathway are more evenly spaced apart when interpreted by the PCI (Figure 1G) compared to the interpretation by the DCI (Figure 1H), which exhibits clusters separated by large distances. This suggests that each method extracts distinct structural insights from glycans.

### The PCI and DCI Segregate Complex N-Glycans According to Structural Themes

As proof of principle that the PCI can segregate the structural information in complex N-glycans in a way that is intuitive and interpretable, N-glycans from chicken colon (Suzuki et al., 2021) were interpreted with the PCI, their results vectorized, and then combined for analysis via PCA. Figure 2 shows that the N-glycans clustered based on structural themes: glycans with multiple bifurcations, glycans with bi- and tri-furcations, glycans with a base node fucosylation, and glycans with differing branch lengths after a bifurcation. The clear aggregation according to structural themes highlights the precision of the PCI for capturing features as subtle as the addition of one residue, a fucose, at the base node, which is well known to have a significant biological effect on the influence exerted by that N-glycan.

**Figure 2.**
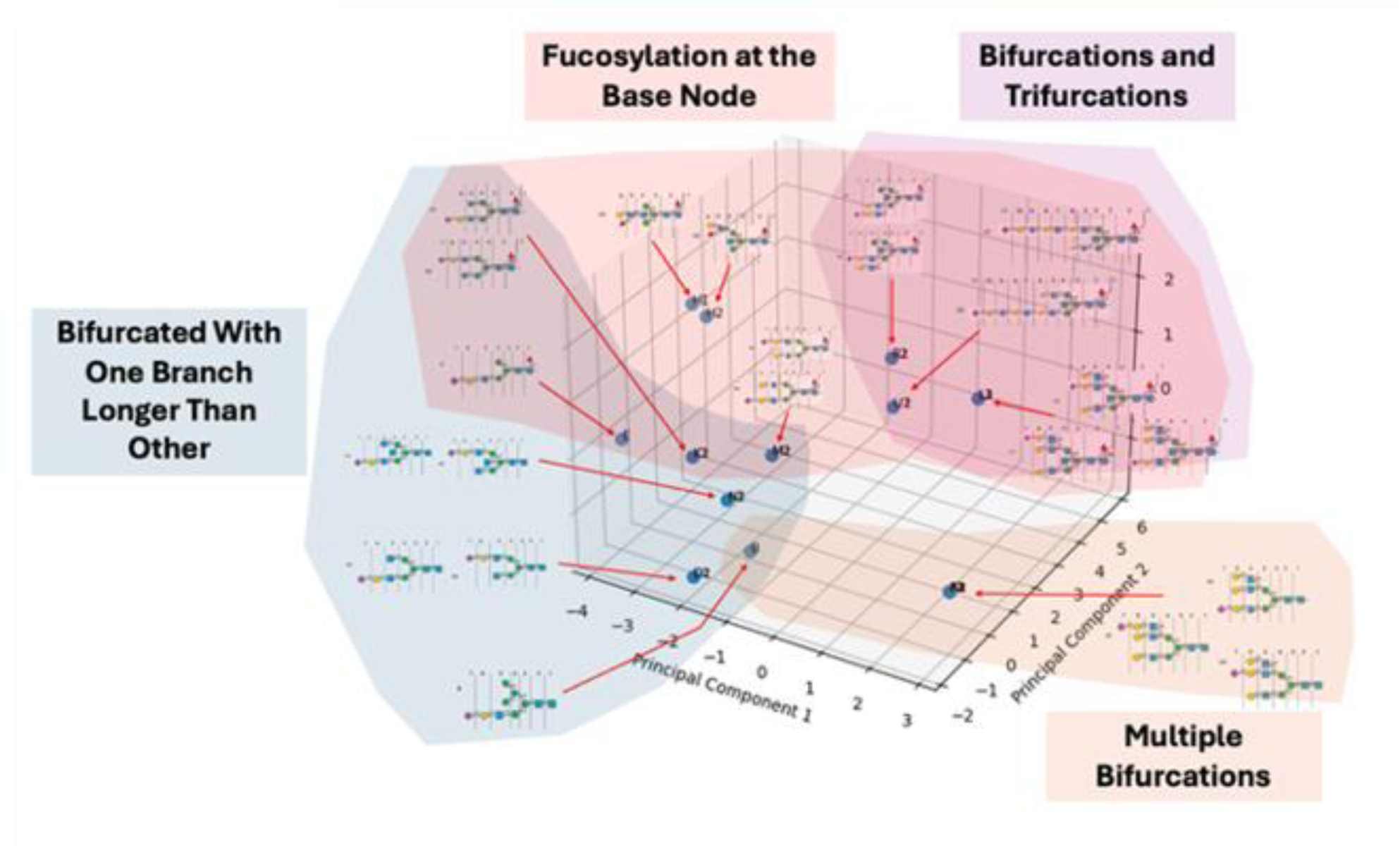
The PCI Segregates N-Glycans from Chicken Colon According to Structural Themes. N-glycans profiled by Suzuki, et al. (2021) were analyzed via the PCI. Their matrices were vectorized as described in the methods, before PCA was performed. The first three principal components were graphed, revealing that the glycans clustered according to structural themes.

The primary goal of this study was to show that the PCI and DCI can extract structural information within N-glycans for insights into prostate cancer. Conroy et al. (2021) measured N-glycans from benign prostate tissue or prostate tumors. PCA on the vectorized DCI matrices reveal that N-glycans that are enriched in prostate tumors cluster according to structural themes (Figure 3) similar to that observed with the chicken colon glycans from Suzuki et al. (2021) in Figure 2. The three major themes are N-glycans with double bifurcation, those with triple bifurcation, and those with one trifurcation. As also observed in Figure 2, the N-glycans in prostate cancer in Figure 3 that had a fucosylation only at the base node associated with higher values on the positive axis of the third principal component. This is consistent with Llop et al.’s (2016) report that alterations in core fucosylation on PSA N-glycans are prime indicators of malignant prostate states.

**Figure 3.**
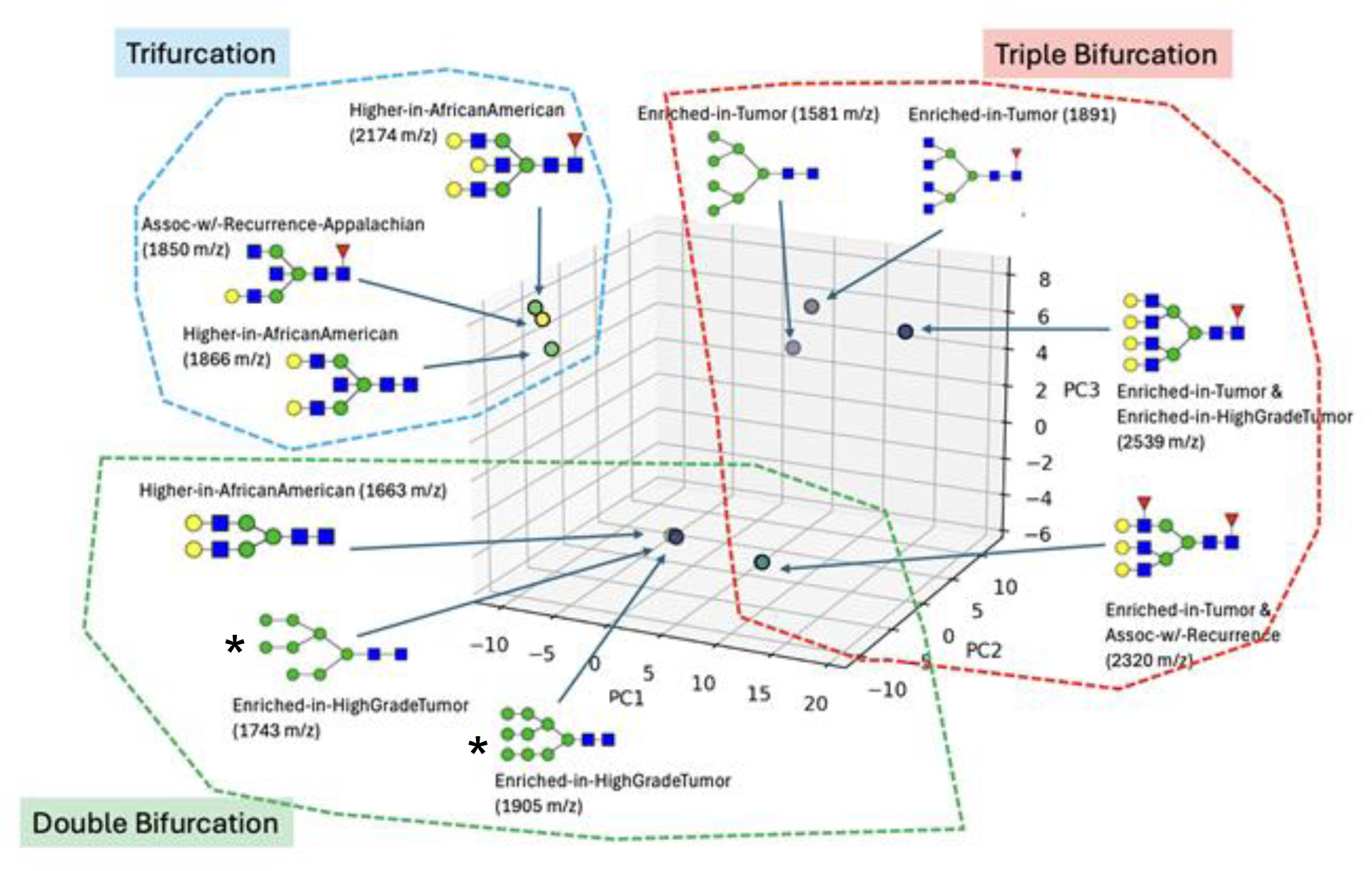
Prostate Cancer N-glycans Segregate According to Structural Themes. Plotting the first three principal components reveal unique clustering according to structural themes: single bifurcations, double bifurcations, and trifurcation. PCA was performed on the vectorized matrices of the PCI for N-glycans from Conroy et al. (2021), which profiled glycans from normal prostate or prostate cancer. *N-glycans that are steps 10 and 11 of the canonical mammalian synthesis pathway (Figure S1).

It is worth noting that two of the structures in the “Double Bifurcation” group (Figure 3, asterisks) are the structures of step 10 and step 11 — whose canonical names are M8 and M9, respectively – in the canonical eukaryotic N-glycan synthesis pathway (Figure S1, Figure 1G&H). The fact that the complex N-glycans associated with prostate tumors cluster distinctly away from these two canonical structures suggests that the 3D distance of “cancerous” glycans from a rationally derived region of “healthy” or “normal” glycans may be useful as a biomarker for cancer in general. Because this distance in the PCA space represents the structural information captured by the DCI, it can serve, at the least, as a quantitative metric for assessing the “amount of information change” in complex, cancer-associated N-glycans relative to a baseline representing glycans associated with “normal” reference. The distance relative to this baseline can also segregate N-glycans from healthy tissue states in the same context as it does for tumor-associated N-glycans. Since this PCA space is 3D, a fuller representation of a glycan’s distance from the canonical pathway should include not just the magnitude of linear 3D distance, but three angles that capture direction. In summary, once a baseline is defined, it is possible to map the distance and direction of all N-glycans in health and disease like star constellations in the night sky relative to Earth. Future work in this direction is warranted for gaining insights into how disease alters N-glycans in ways not detected by m/z ratio and relative abundance, but that are complementary to them.

The second set prostate cancer glycans analyzed in this study were from Tajiri et al. (2007) wherein glycans attached to PSA from prostate cancer patients were extracted. The N-glycans were either from PSA in plasma of seminal fluid or from PSA in serum plasma. In the case of PSA from serum plasma, the PSA was either free or attached to α1-antichymotrypsin (ACT). Figure 4 plots the first three principal components of the vectorized DCI matrices, revealing that all the N-glycans from seminal fluid PSA occupy a distinct 3D region (red dotted oval), while N-glycans from serum PSA, whether free or bound to ACT, can exist either in this region or outside of it. The most striking observation in this data is that some glycans from serum PSA are more similar to glycans from seminal fluid PSA compared to other glycans are also from serum PSA. 25-40% of N-glycans from serum PSA are located far outside the red dotted region representing seminal plasma, which is an objective and quantitative stratification. The results suggest the existence of structural subtypes, as defined by spatial information captured by the DCI: in this case, “seminal fluid-like” and “non-seminal fluid-like.” Biochemically, this distinct “non-seminal-like” structural domain captured by the DCI can potentially correlate with the documented variations in core fucosylation and specific sialylation patterns characteristic of serum PSA in malignant states (Llop et al., 2016). Such structural subtypes are not identifiable by traditional metrics, such as m/z ratio, as shown in Figure 4B wherein the m/z ratios obtained from Tajiri et al. (2007) are plotted. In Figure 4B, while all serum PSA glycans that are inside the red dotted oval of Figure 4A have m/z ratios that are within the range of seminal plasma PSA (meaning below the blue dotted line), m/z ratio alone provides no reason to suspect that there should be a “seminal fluid-like” subtype. A m/z ratio does not explain the type of branches or number of branches in a glycan, meaning multiple different glycan structures can have similar m/z ratios. It is the PCA space of the DCI matrices that suggest the existence of two types of serum PSA glycans with reference to seminal fluid PSA glycans.

**Figure 4.**
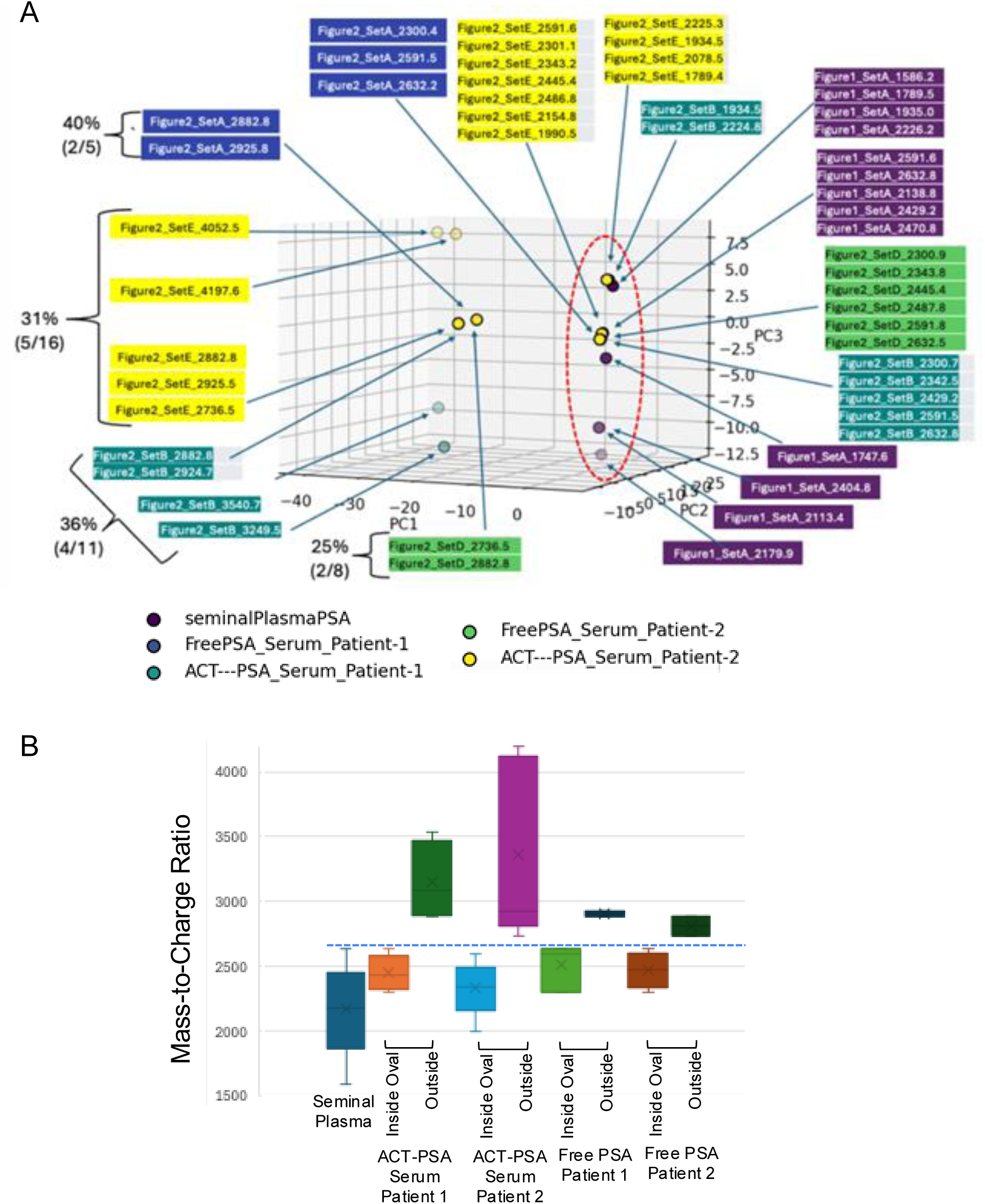
**N-glycan Structural Information Identifies Serum Plasma Glycans as Being Seminal Plasma-Like vs. Serum Plasma-Like** (A) Tajiri, et al. (2007) profiled N-glycans attached to the PSA protein from either seminal fluid or serum. The first three principal components of vectorized DCI matrices were plotted. N-glycans from seminal plasma contain spatial information that occupies a distinct region of 3D PCA space (red dotted oval). 25% to 40% of glycans from serum plasma PSA are spatially distinct from seminal plasma, which is objectively stratified by this approach. (B) The m/z ratios of glycans in panel A are grouped according to whether they are in the red dotted oval in panel A, which represents “seminal fluid-like” glycans, or outside of it. The blue dotted line represents the maximum value for the seminal plasma group.

### Stratifying Prostate Cancer Stages as a Function of Glycan Structural Information

Nyalwidhe et al. (2013) profiled N-glycans from expressed prostatic secretions (EPS) of patients that were at various stages of prostate cancer: non-cancer, indolent cancer, and aggressive cancer. N-glycans from EPS urine or exosomes from EPS were profiled at each of these three stages. Figure 5A shows the first three principal components of PCA on vectorized DCI matrices of N-glycans that were categorized as either being present in both cancer and non-cancer conditions or that are unique to any cancer condition. Notably, glycans that are present in both cancer and non-cancer conditions occupy a narrower 3D space (purple dotted box) than glycans that are unique to any cancer condition (yellow dotted box). This finding is in accord with the understanding that cancerous cells, when they dedifferentiate, shed restrictions that apply to non-cancer cells. From a biochemical perspective, this expanded structural space likely reflects the loss of enzymatic fidelity and the subsequent biosynthetic hyper-heterogeneity of glycan processing typical of malignant transformation (Pinho & Reis, 2015).

**Figure 5.**
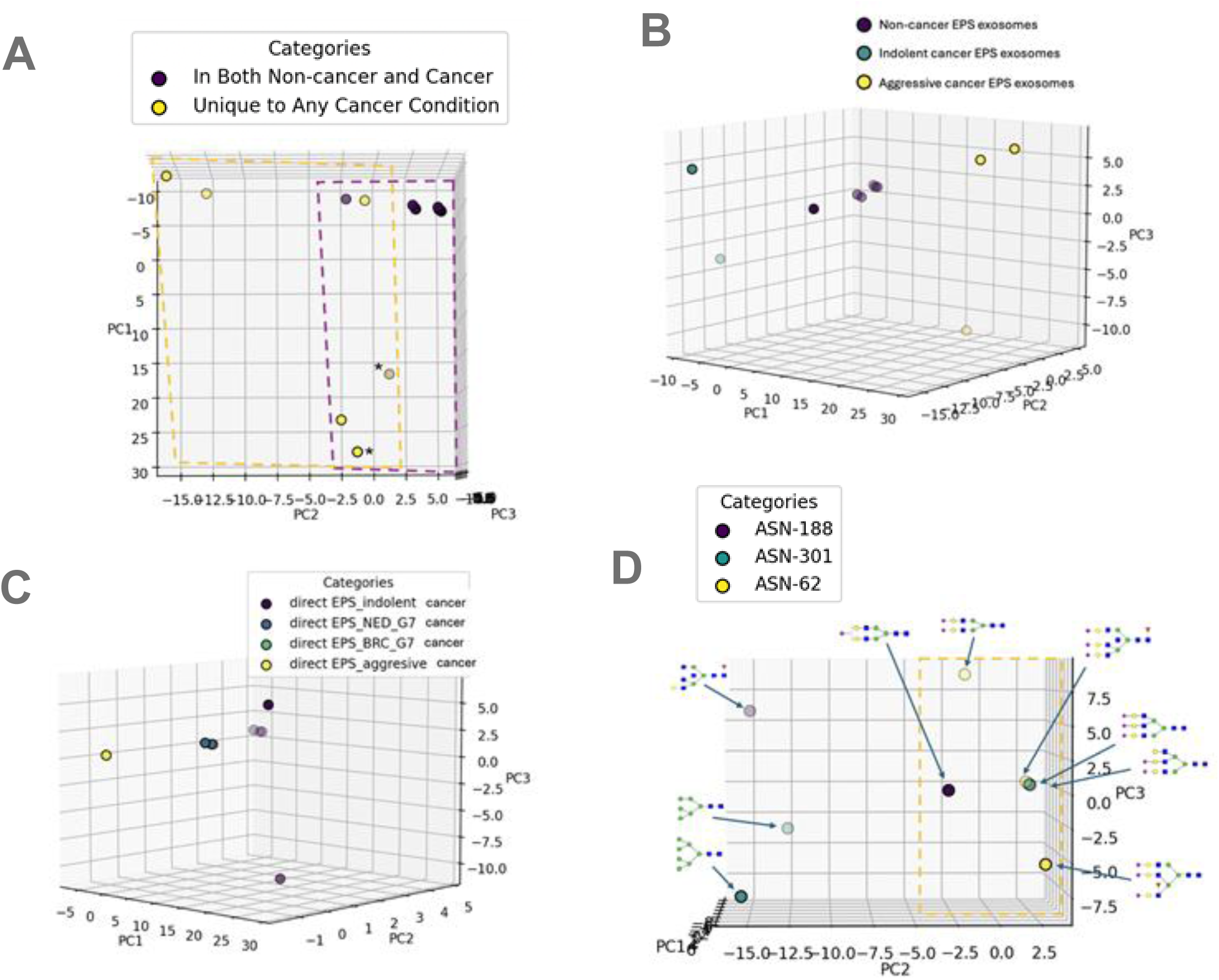
Segregation of N-glycans as a Function of Prostate Cancer Stage Nyalwidhe, et al. (2013) profiled N-glycans from EPS of patients that were at various stages of prostate cancer. (A) The first three principal components of vectorized DCI matrices were plotted for N-glycans that were either present in both non-cancer and cancer conditions vs. those that were unique to any cancer condition. *This glycan is in both non-cancer and cancer conditions in one of the three EPS sources. (B) The same analysis as in A, but for N-glycans located on exosomes from EPS from non-cancer cases, indolent cancer cases, or aggressive cancer cases. (C) The same analysis as in A, but for N-glycans from EPS of indolent cancer, Gleason 7 cancer with no evidence of disease (NED) in 3 years or greater postprostatectomy, Gleason 7 cancer with biochemical recurrence (BRC) within 6 months of prostatectomy, or aggressive cancer. (D) Same analysis as in A, but for N-glycans attached to specific glycosites on the PAP protein extracted from aggressive prostate cancers.

When N-glycans from either exosomes in EPS (Figure 5B) or EPS urine (Figure 5C) were analyzed by the DCI, a clear stratification showing a progression from non-cancer to indolent cancer to aggressive cancer is observed. The distinct clustering of glycans according to cancer stage, separated by large regions of 3D space, suggest that the vectorized DCI can objectively and quantitatively stratify the effect of prostate cancer on N-glycan structural information.

Clinical studies like McKiernan et al. (2018) have already established a validated clinical benchmark for using EPS urine exosomes in non-invasive prostate cancer risk stratification to avoid unnecessary tissue biopsies. While their commercial assay (ExoDx) relies on gene expression, the results of this study can prove that N-glycan structural information from the exact same clinical matrices (EPS exosomes and urine) provides a robust, independent informational layer for stage stratification. This positions GlycoAssessor as a valuable tool for glycan-based liquid biopsies. These results demonstrate the robust translational utility of the DCI framework to capture subtle glycoarchitectural changes within a clinical context, highlighting its potential for non-invasive liquid biopsy development in alignment with current exosome-based diagnostic trends (McKiernan et al., 2018).

### Glycosite-Specific Insights From a Glycosylated Cancer-Associated Protein

Nyalwidhe et al. (2013) also profiled the N-glycans attached to Prostatic Acid Phosphatase (PAP), a prostate cancer biomarker, from the EPS of aggressive prostate cancers. PAP has three glycosites located at asparagine (Asn) residues: Asn-62, Asn-188, and Asn-301. Figure 5D shows the first three principal components of PCA done on vectorized DCI matrices of N-glycans attached to these three glycosites, revealing a spatial distribution in PCA space that is unique to Asn-62. Nyalwidhe et al. (2013) report that “As with the seminal fluid PAP sample, the third site at Asn 301 is the only site where high-mannose structures are determined,” which highlights the fact that there are natural restrictions on what type of N-glycans can be attached to specific glycosites on PAP. This knowledge of restrictions on what can be attached at each site suggests that Asn-62 has a narrow band of structural information (defined as PCA space) that can be attached to it (Figure 5D, yellow dotted box) compared to Asn-188 and Asn-301, which can harbor glycans with more diverse structural information.

We hypothesize that the narrow band observed for Asn-62 reflects local physical or steric constraints, whereas the permissive nature of Asn-188 and Asn-301 suggests they reside in more flexible, solvent-exposed regions. This biological explanation is strongly supported by the 3D crystal structure of human PAP published by Jakob et al. (2000), which reported that the nearby residue Glu-63 experiences a massive conformational shift because physical forces shift the carbohydrate chain at Asn-62 in a tight direction (p. 213). Due to these structural limitations, Asn-62 is restricted exclusively to smaller, rigid high-mannose chains (p. 211, 216). Conversely, Asn-188 is capable of accommodating bulky, highly branched complex oligosaccharides (specifically described as the longest chain with bi- or triantennary structures and fucose attachments) because it has more space (p. 216).

Restrictions and permissiveness of allowable structures at specific glycosites have testable implications about the role of glycosites in the function of PAP in prostate cancer. Because the spatial information, defined as PCA space, allowed at Asn-62 is more limited, this can be interpreted to mean that this site has a lower tolerance for changes within glycan structure. This restriction has important implications for therapeutic modalities that attach chemical moieties to glycans, whether to de-activate, activate, or modulate the associated protein’s function. On the other hand, the fact that Asn-188 and Asn-301 have a broader range of allowable spatial information means that adding chemical moieties to glycans at these sites may result in a desired modification to the protein’s behavior without triggering drastic deactivation. Regardless of the interpretations, they are empirically testable.

## Discussion

The central vision of this study is to demonstrate that spatial information encoded within N-glycan structures constitutes a biologically meaningful and quantifiable metric that can be leveraged to stratify prostate cancer states. This work seeks to validate the DCI and the PCI as robust computational frameworks for classifying structural themes within N-glycans and for distinguishing healthy from diseased states based on glycan monomer spatial organization rather than chemical composition. A key objective is to establish the GlycoAssessor software as a reliable, open-source, and user-friendly software platform capable of calculating DCI and PCI from glycans represented in SNFG standards, while also providing accessible educational resources to facilitate broader adoption of these methods. Other methods to classify and identify glycan structures exist such as glypy (Klein & Zaia, 2019) and GlycanCT (Akihiro et al., 2026), though their scopes differ significantly. Using prostate cancer–associated N-glycan datasets, this study aims to demonstrate that DCI and PCI metrics can stratify benign and malignant tissue states and further resolve disease progression across stages through unsupervised dimensionality reduction. By extending these analyses across multiple datasets, this work proves that consistent spatial glycan patterns persist despite biological and experimental heterogeneity.

### Assumptions and Limitations

An assumption underlying the DCI and PCI algorithms is that monosaccharide identity is not required to capture disease-relevant structural information. By focusing on spatial arrangement, these methods assume that branching architecture and connectivity encode sufficient biological signals. While this enhances generalizability, it may overlook composition-specific effects that contribute to function. However, it is possible to incorporate the importance of monosaccharide identity into these metrics if a mapping system is created that adds weights to the values within the DCI and PCI matrices. For example: The English alphabet has 26 letters that are ordered from a to z, and a rudimentary mapping system could assign the value of 1 to a, 2 to b, 3 to c, etc. For monosaccharides, however, there needs to be a more complex mapping system to capture the significance of their weights because there are recurring themes that dictate monomer locations (e.g. eukaryotic N-glycans always begin with two GlcNAc; fucosylations only occur with fucose). One way to derive a meaningful mapping system for humans would be to examine all known human N-glycans that occur in healthy states, and then calculate the frequency at which they occur in each layer of a glycan, as defined by the PCI (Figure 1A). This frequency for any type of monosaccharide, such as mannose, can be incorporated into the PCI matrix or DCI matrix as a weight that is multiplied into specific features or incorporated as additional columns. How to exactly incorporate this information needs to be optimized and is beyond the scope of this study. The approach just described is similar to the method called “frequency encoding” that is used in data science to turn categorical variables into quantitative variables (Pargent et al., 2022), except that a sophisticated, and biologically meaningful, mapping system is required for N-glycans. This approach of developing a mapping system for specific monosaccharides applies to more than just humans. For example, this is applicable in viral, prokaryotic, eukaryotic, mammals, or other glycoproteins. In each case, knowing all N-glycans is not feasible and every group has their own specificities. Therefore, a consensus could be found for different groups in the future.

On a separate note, it is important to note that the vectorization process for the PCI and DCI creates a vector space that is not absolute, meaning the length and composition of the vector is largely, but not solely, determined by the most complex N-glycan being compared in a group of N-glycans. However, this means that if all possible N-glycans for a type of cancer can be known, then it is possible to create a “universal vector space” for that cancer. This can be done by vectorizing all known N-glycans in that group according to the unique features present. In other words, any N-glycan would be vectorized based on the features of a composite, imaginary N-glycan that contains all features within a group. This same approach can be applied to any group of organisms or cell types. However, for organisms like yeast, which can have multiple recurring branching units that are attached in tandem, it may be difficult to derive universal vector space. In this case, a rationale is needed as to why a “quasi-universal” space is acceptable.

---

**Figure S1.**
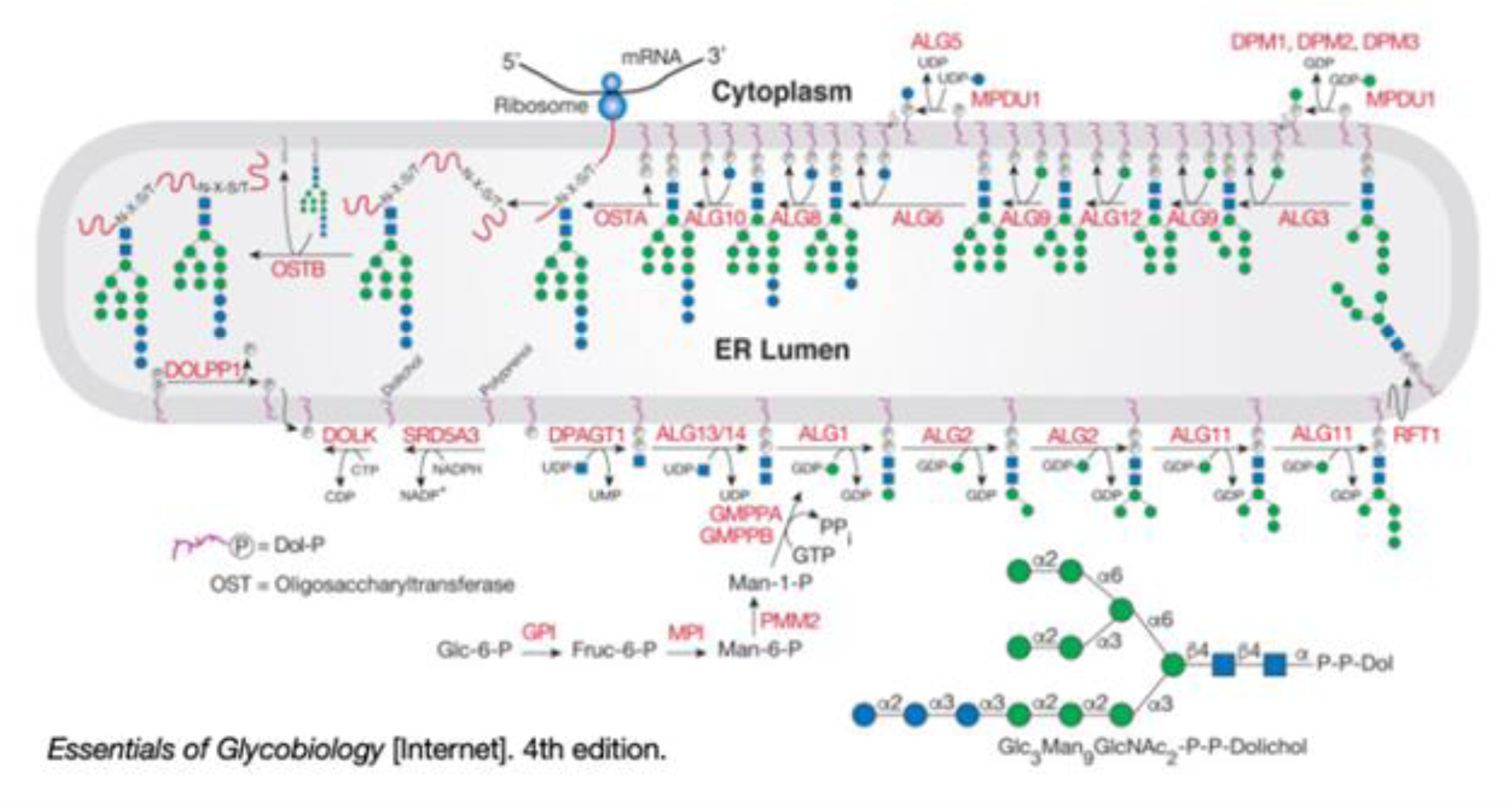
Pathway for Synthesis of N-glycans in Eukaryotes Image source is Stanley, et al. (2022). Shown is synthesis of dolichol located on the cytoplasmic face of the ER. In mammalian cells, the oligosaccharyltransferase-A (OST-A) complex is associated with the translocon in the ER membrane and preferentially glycosylates nascent polypeptides traversing the translocon, whereas the OST-B complex modifies proteins that have left the translocon and are in the ER lumen.

